# Analysis of structural variants in four African Cichlids highlights an association with developmental and immune related genes

**DOI:** 10.1101/473710

**Authors:** Luca Penso-Dolfin, Angela Man, Wilfried Haerty, Federica Di Palma

## Abstract

African Lakes Cichlids are one of the most impressive example of adaptive radiation. Independently in Lake Victoria, Tanganyika, and Malawi, several hundreds of species arose within the last 10 million to 100,000 years. Whereas most analyses in Cichlids focused on nucleotide substitutions across species to investigate the genetic bases of this explosive radiation, to date, no study has investigated the contribution of structural variants (SVs) to speciation events (through a reduction of gene flow) and adaptation to different ecological niches. Here, we annotate and characterize the repertoires and evolutionary potential of different SV classes (deletion, duplication, inversion, insertions and translocations) in five Cichlid species (*Haplochromis burtoni, Metriaclima zebra, Neolamprologus brichardi, Pundamilia nyererei* and *Oreochromis niloticus*). We investigate the patterns of gain/loss across the phylogeny for each SV type enabling the identification of both lineage specific events and a set of conserved SVs, common to all four species in the radiation. Both deletion and inversion events show a significant overlap with SINE elements, while inversions additionally show a limited, but significant association with DNA transposons. Inverted regions are enriched for genes regulating behaviour, or involved in skeletal and visual system development. We also find that duplicated regions show enrichment for genes associated with “antigen processing and presentation” (GO:0019882) and other immune related categories. Altogether, we provide the first, comprehensive overview of rearrangement evolution in East African Cichlids, and some initial insights into their possible contribution to adaptation.

## Introduction

African Cichlids represent one of the best examples of rapid adaptive radiation (Kocher 2004; Wagner, et al. 2012). The adaptation to different ecological niches in Lakes Malawi, Tanganyika and Victoria has given rise to several hundreds of species in a period of just a few million years. The radiation is associated with great phenotypic variation, including jaw morphology, body shape, coloration, adaptation of the visual system to different water depths, and behaviour.

Variation in ecological niches and behaviour appears to be associated with different brain development (Huber, et al. 1997), with differences appearing already at early developmental stages (Sylvester, et al. 2013).A great example of adaptation is represented by the evolution of the Cichlid visual system, involving eight different opsin genes (Fernald 1984; Hofmann, et al. 2009; Muschick, et al. 2011).

To gain insights on the molecular mechanisms underlying this rapid radiation, Brawand *et al.* (Brawand, et al. 2014) generated genome references for five species: the Nile tilapia *(Oreochromis niloticus)*, representing an ancestral lineage; *Neolamprologus brichardi* (Lake Tanganyika), *Metriaclima zebra* (Lake Malawi), *Pundamilia nyererei* (Lake Victoria), and *Haplochromis burtoni* (riverine species around Lake Tanganyika). The study highlighted several mechanisms underlying species diversification, including selection acting on existing standing variation, high rates of gene duplication, novel microRNAs and rapid sequence divergence in otherwise conserved non-coding elements. Following this study, Malinksy *et al.* described an example of early stage divergence between two Cichlid ecomorphs in Tanzania. (Malinsky, et al. 2015). They identified genomic islands of speciation between them, containing potentially adaptive genes associated with mate choice. Theis *et al.* (Theis, et al. 2014) focused on the early phases of adaptive divergence of *H. burtoni,* which is found in both Lake Tanganyika and inflowing river. Their results highlighted the presence of multiple divergent lake-stream populations, representing different stages of the speciation process. More recently, the sequencing of 134 individuals covering 73 species provided a great characterisation of genomic diversity in lake Malawi (Malinsky, et al. 2017). The authors observed very low levels inter-species divergence (0.1-0.25%), overlapping the diversity found within species. Phylogenetic analyses showed that no single species tree can efficiently represent all species relationships, suggesting high levels of repeatedly occurring gene flow.

All studies so far focused on single variation within and between species and to a lesser extent on the evolution of gene regulatory patterns (Mehta *et al.*, in preparation). To the best of our knowledge, the role of structural variation in the evolution of East African Cichlids has not been investigated yet. Structural variants (SVs, including deletions, duplications, inversions, insertions and translocations) are the source of increased genomic variability and in some cases adaptive potential. Gene duplication events might lead to neo-or sub-functionalisation of the resulting paralogues (Ohno 1970; Lynch 2002; Katju and Lynch 2006), while deletion can reflect relaxed selective pressure or be possibly adaptive in other cases (Sharma, et al. 2018). Inversions result in supressed recombination when heterozygous, and might act as a protection against gene flow for specific haplotypes (McGaugh and Noor 2012). Inversions might raise in frequency, up to fixation, possibly leading to isolation and even speciation events (Kirkpatrick 2010; Catacchio, et al. 2018). Studies in *Drosophila melanogaster* provided strong evidence for their involvement in adaptation. For example, the inversion **3RP** is associated with adaptation to different climates (Rane, et al. 2015). Its frequency exhibits a parallel latitudinal cline across several continents, being higher close to the equator and decreasing towards higher latitudes (Kirkpatrick 2010; Rane, et al. 2015).

Translocations can result in a heavy restructuration of chromosome organisation (Rowley 2001), with potential gene loss or changes in regulatory control of expression.

Identifying structural changes across species representative of all three great lakes can provide exciting insights into their explosive radiation. In this study, we use the newly released *O. niloticus* reference based on long read PACBIO sequencing (Conte, et al. 2017) and paired-end sequencing data generated by (Brawand, et al. 2014) to identify SVs in four Cichlid species, representative of Lake Malawi, Victoria and Tanganyika. Through this analysis we aim to: characterise the evolutionary patterns associated with different rearrangement classes, investigate functional enrichment within those rearranged genomic regions, and to identify the genes affected by these structural changes and how these can relate to the phenotypes found across the three lakes.

We show that genes lying inside inverted regions are enriched for genes regulating behaviour, or involved in skeletal and visual system development, which are directly relevant to the African radiation. Altogether, we describe the repertoires of structural variations across four species of the East African Cichlids, their evolutionary dynamics, and novel insights into their possible contribution to adaptation.

## Results

### Annotation of SVs across 4 Cichlid species

We mapped all previously generated paired-end libraries (Brawand et al. 2004) to the high quality *O. niloticus* assembly (Conte, et al. 2017) to annotate five different classes of rearrangements (deletion, tandem duplication, inversion, insertion and translocation) in the four available species of the East African radiation (Supplementary Fig. 1). We used a combination of three different tools: *Breakdancer* (Fan, et al. 2014), *Delly* (Rausch, et al. 2012) and *Pindel* (Ye, et al. 2009) and identified 6694 deletions, 1550 duplications, 1471 inversions, 34875 insertions and 1354 translocations (table 1).

**Table 1.** Annotation of SVs

| SV class | Total | >1 species | All species |
| --- | --- | --- | --- |
| DEL | 6694 | 3903 | 541 |
| DUP | 1550 | 353 | 22 |
| INV | 1471 | 566 | 188 |
| INS | 34875 | 18253 | 3949 |
| TRA | 1354 | 92 | 2 |

Our initial predictions showed a bias towards small (<1kb) deletions (240229). This number might be inflated as a result of our SV detection pipeline, where deletions are identified using read pairs mapped in a concordant way (as opposed to duplications and inversions). This represents an issue particularly when considering small events. Therefore, we decided to retain only deletions with a minimum size of 1kb (Table 1). In the resulting dataset, 5483 deletions fall in the 1-10kb size range, while 1207 represent larger, >10kb events (Fig. 1). We investigated whether the size of a SV correlates with the age of the event. While the size distributions of deletions did not seem to be affected by the number of species sharing the SV, we noticed a tendency for duplications and inversions towards larger sizes as the number of species increased. Species specific events are significantly smaller than those common to 2 species (MW test, p=0.005), which in turn are smaller than the events found in 3 species (MW test, Table S1). Moreover, conserved inversions are significantly larger than both conserved deletions (MW test, p<0.0001) and duplications (MW test, p<0.0001).

**Fig. 1.**
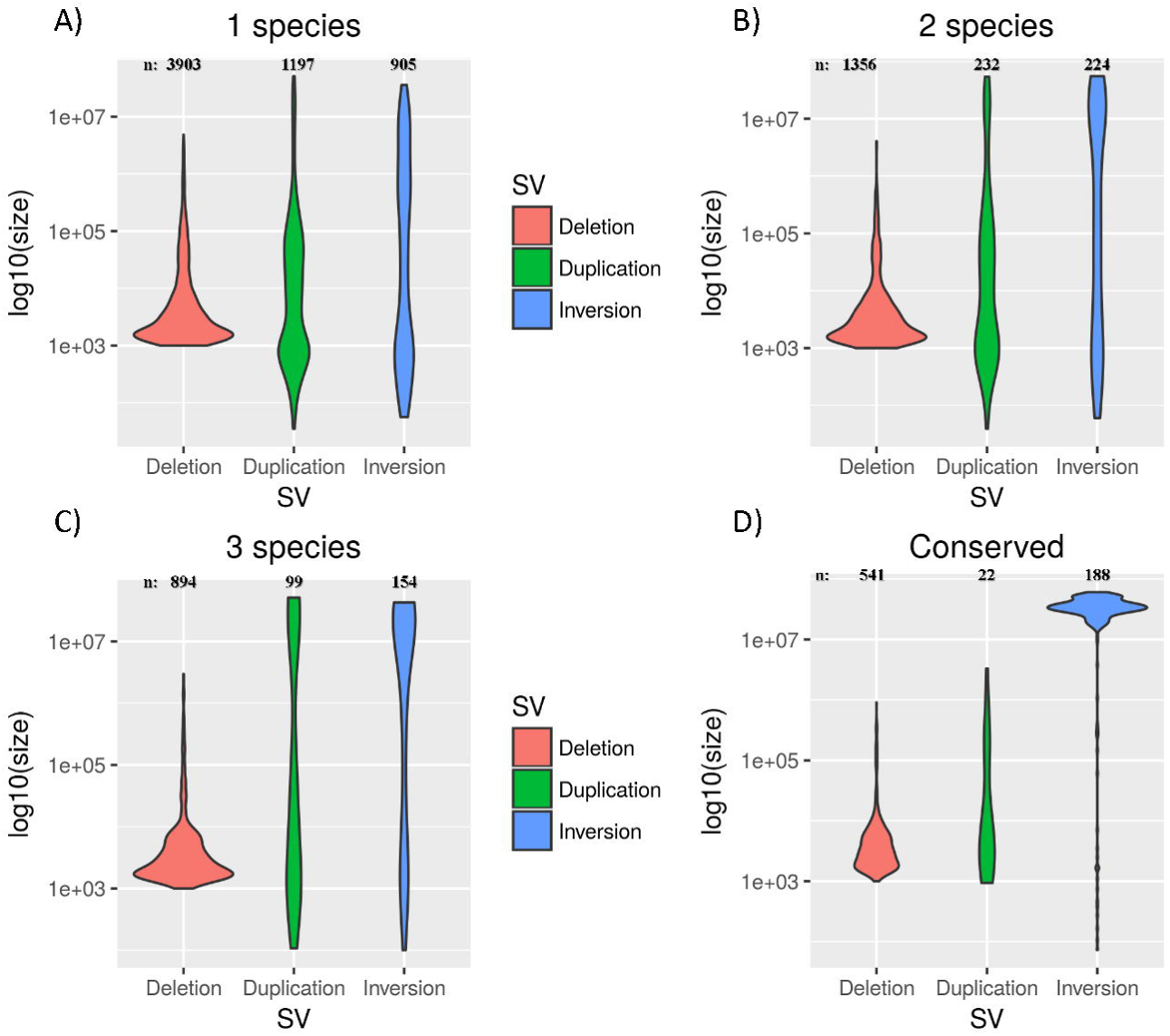
Violin plots of the size of different intra-chromosomal SV classes, considering different levels of conservation (*conserved* refers to SVs common to all 4 species).

We investigated the patterns of gain and loss evolution for each SV class, using a *Dollo parsimony* approach (see Materials and Methods). We identified a high proportion of events predicted to be lineage specific (Fig.2). Additionally, comparison across species allowed us to identify the events common to a single lineage or to all species involved in the African radiation. We will refer to the latter as “conserved SVs”.

**Fig. 2.**
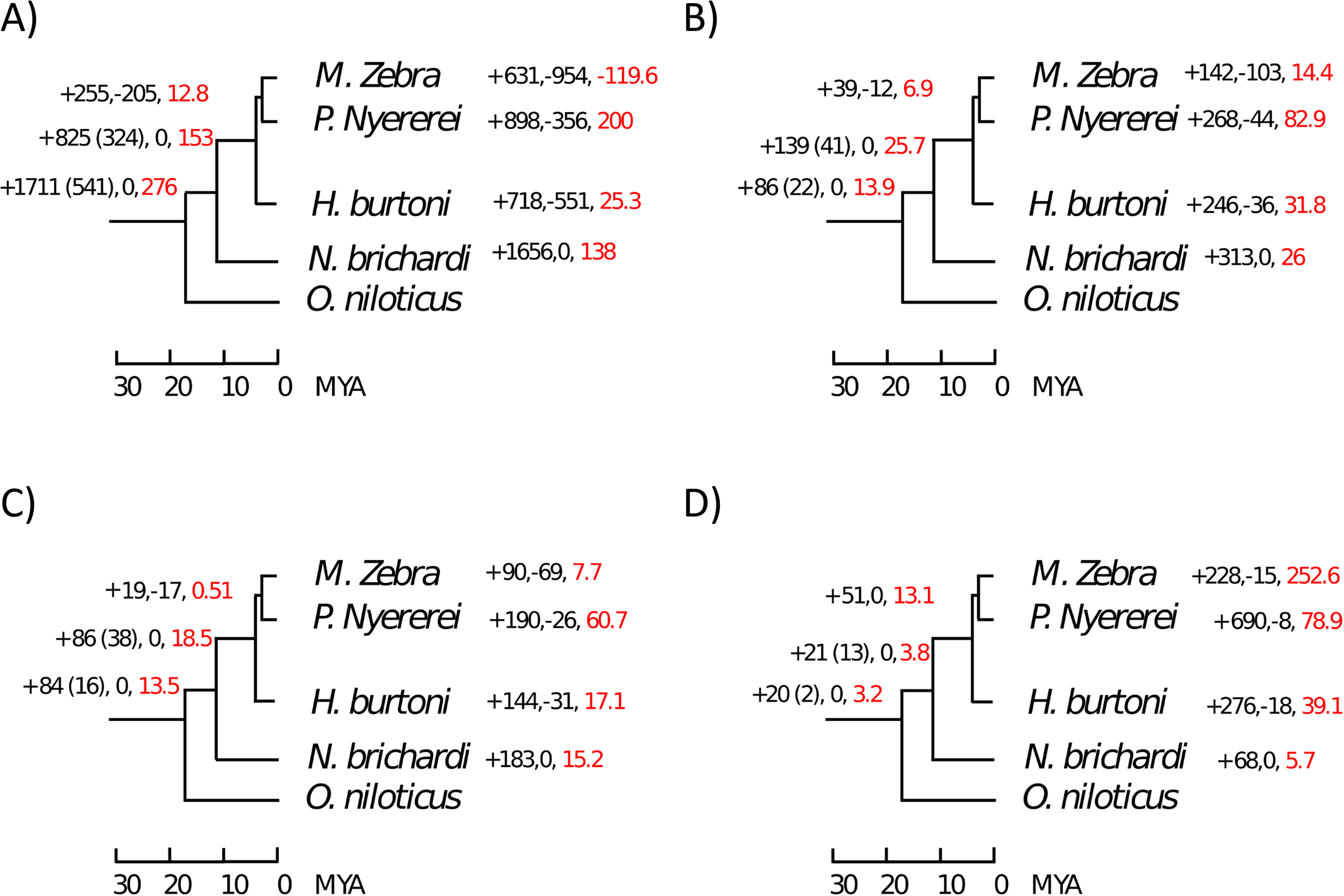
Gain and loss of different SV types (up to 5Mb in size) across the phylogenetic tree. For each branch, the number of gained and lost events is provided, as well as the net gain rate per million years (red labelled). When different from the total number of gains, the number of events which are gained and retained across the whole lineage (not lost afterwards) is indicated in brackets.

We noticed a surprisingly high loss rate of deletions in the *M. zebra* lineage (Fig.2A). In order to evaluate the reliability of our approach and the accuracy of our annotations, we compared our results to the those obtained through the pairwise, whole genome alignments between the latest *M. zebra* and *O. niloticus* assemblies, using *Satsuma2* (https://github.com/bioinfologics/satsuma2) (see Materials and methods). Out of 2263 deletions annotated in *M. zebra*, only 54 (2%) were discordant with *Satsuma2* alignments.. Thus, we show that our annotation in *M. zebra* has a very high concordance with the high quality genome assemblies of *M. zebra* and *O. niloticus*.

With the exception of *M. zebra* deletions, we observed high proportions of lineage specific events, consistent across all SV classes (Fig.2). However, in the case of deletions we also observed a high number (1711) of events which are ancestral to the radiation. Overall, these results point at a reduction in genome size associated to the African radiation. This is in concordance with the observation that the *M. zebra* assembly is 48Mb shorter than the *O. niloticus* reference (www.ncbi.nlm.nih.gov/genome/genomes/2640;www.ncbi.nlm.nih.gov/genome/genomes/197). However, it must be noted that, in the absence of an outgroup for the five species, these deletions we cannot rule out the possibility that some of these deletions have rather occurred in the *O. niloticus* lineage.

We investigated the extent of interval overlap between our predicted SVs and different genomic features.

We considered different subsets of our SV annotations, categorising our predictions based on size range (<1kb, 1kb-10kb,>10kb) and number of species sharing the SV event (whole dataset vs conserved SVs). We observed a strong association between >10kb conserved deletions and immunoglobulin chain regions. The association is highly significant for both the constant (14.8 fold change, p=0.01) and variable (10.8 fold change, p=5e-03) gene segment annotation, which suggests a possible involvement of copy number variants in immune response mechanisms. It must be pointed out, nevertheless, that these loci are present in multiple, tandemly repeated copies, and the observed association could possibly reflect assembly issues in repetitive regions.

We also hypothesised that repeats throughout the genome facilitate the evolution of structural changes. In order to test this hypothesis, we looked at the genomic association (interval overlap analysis, see Materials and Methods) between our SV dataset and African Cichlid specific repetitive elements. These analyses highlighted a significant overlap between 1kb-10kb long conserved inversions and SINE2 elements (10.2 fold change, p=8.7e-03). The association with SINE2 is not significant, however, when we consider all conserved inversions, irrespective of their size (1 fold change, p=0.3). We identified repeat elements contained inside conserved inversions, and considered their percentage of divergence from the consensus sequence (Fig. 7). We observed high representation, in terms of total length, of L2 elements, especially when copies with 1-4% divergence from the consensus are considered. SINE elements become more represented in higher divergence bins.

Conserved duplications are significantly under-represented with African Cichlids (AFC) SINE2-1 (0.64 fold change, p=2.7e-03) and REX1-2 AFC elements (0.3 fold change, p=3.14e-02). Conversely, they appear to be enriched for several simple repeats, including (AAGTCTC)n (54.7 fold change,p=1e-04).

Large deletions appear to be negatively associated with AFC RTE-2 elements (0.57 fold change, p=3.7e-02) but positively associated with AFC L1-1 elements (2.17 fold change, p=8.7e-03), as well as several simple repeats. When we considered all conserved deletions, irrespective of their size, we observed a significant association with AFC SINE2-1 (1.42 fold change, p=1e-04) and SINE3 (2.98 fold change, p=1e-04) elements. Similar conclusions were reached in previous studies on the pig genome (Ai, et al. 2015; Zhao, et al. 2016). Taken together, these results suggest a correlation between repetitive elements and structural evolution in African Cichlids.

We next asked the question whether we observe differences in the repeat landscape inside and outside SV regions. Previous studies have highlighted very high proportions of DNA transposable elements in African Cichlids (Brawand, et al. 2014). Our data confirms these findings, as shown in figg 3-5.

**Fig. 3.**
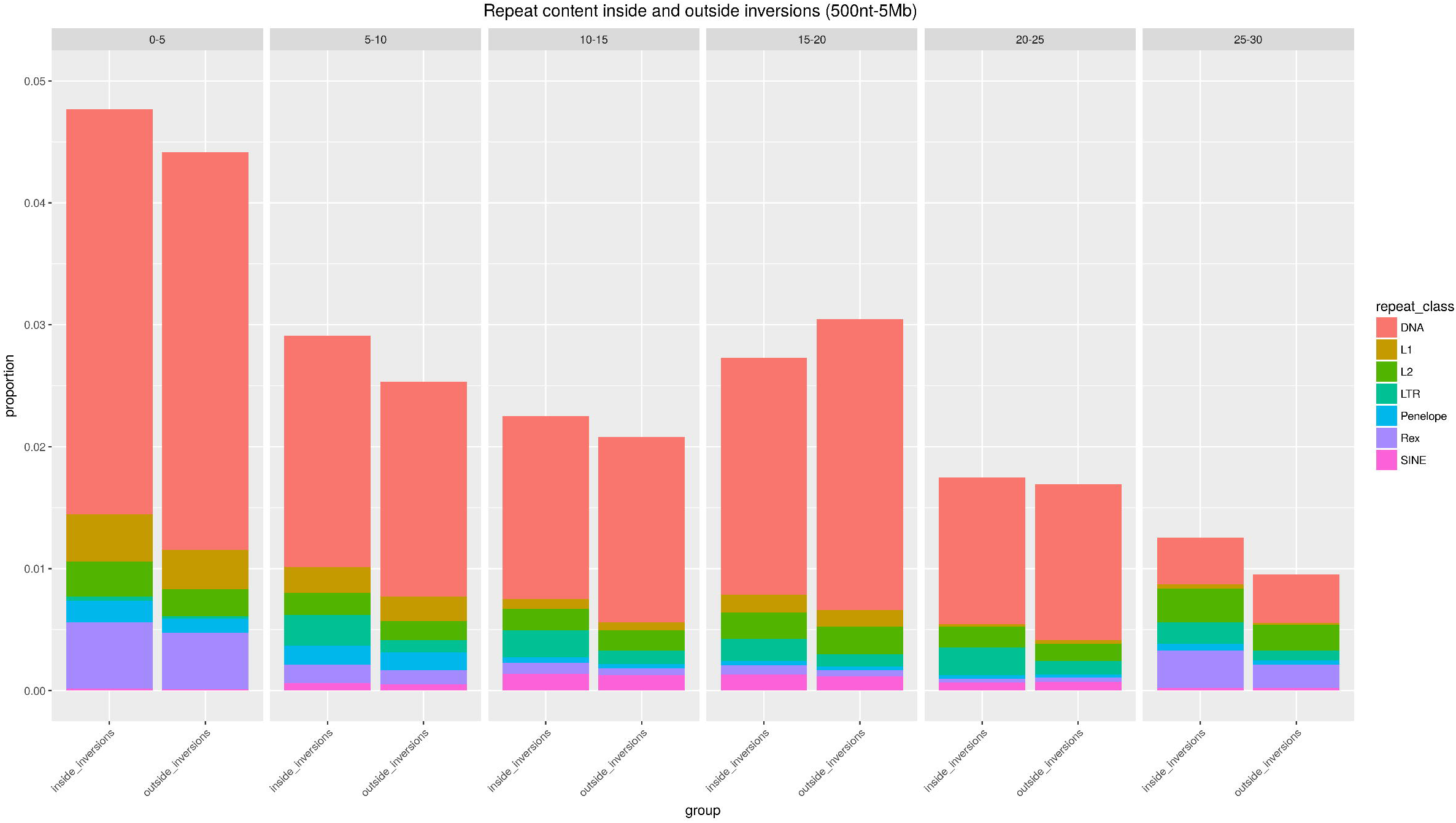
Proportion of nucleotides inside and outside *M. zebra* inversions which are part of a repeat element, grouped based on the percentage of divergence from the consensus. Different colours correspond to distinct repeat classes. Each grid corresponds to a specific divergence interval.

**Fig. 4.**
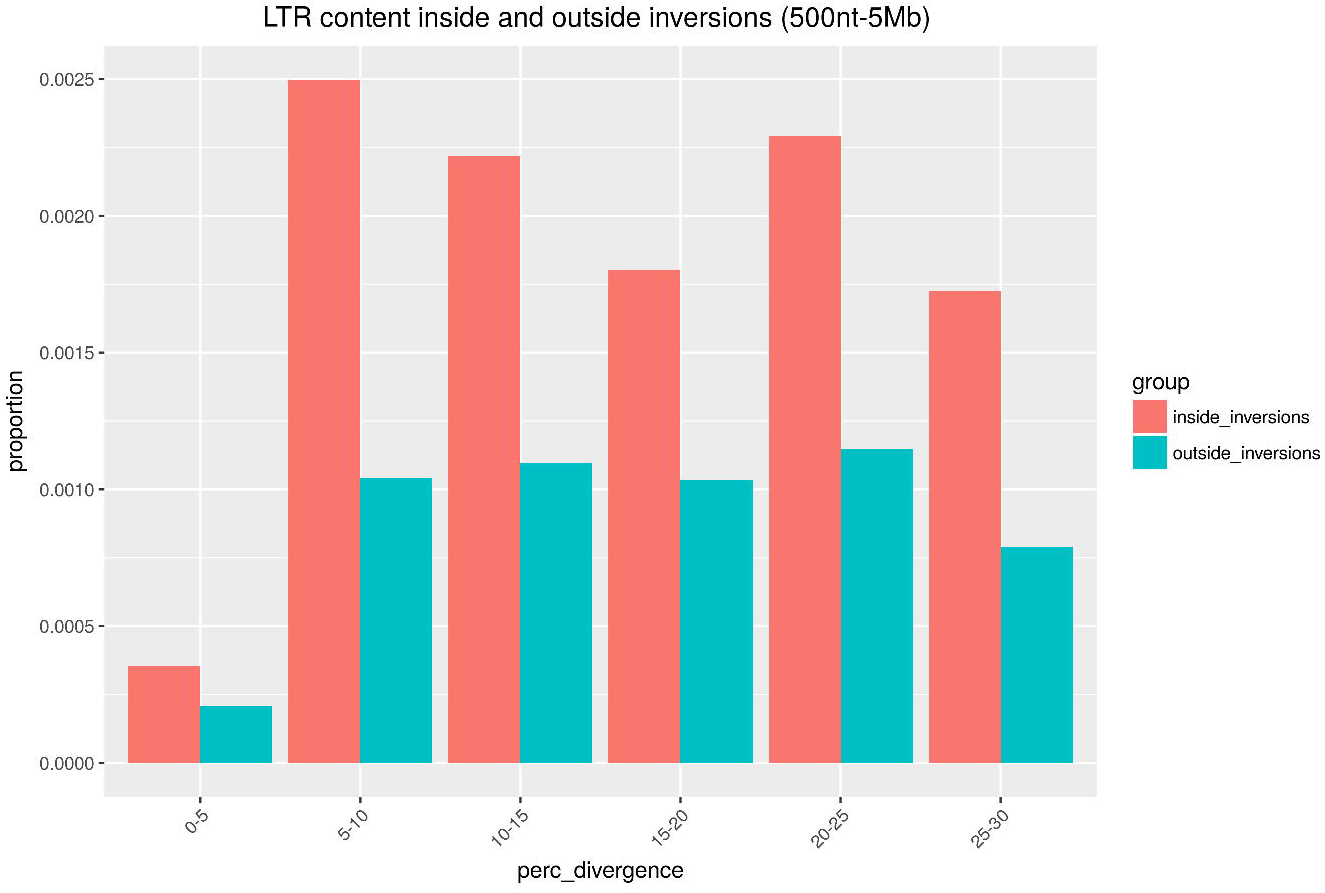
Proportion of nucleotides inside and outside *M. zebra* inversions which are part of an LTR elements, grouped based on the percentage of divergence from the consensus.

**Fig. 5.**
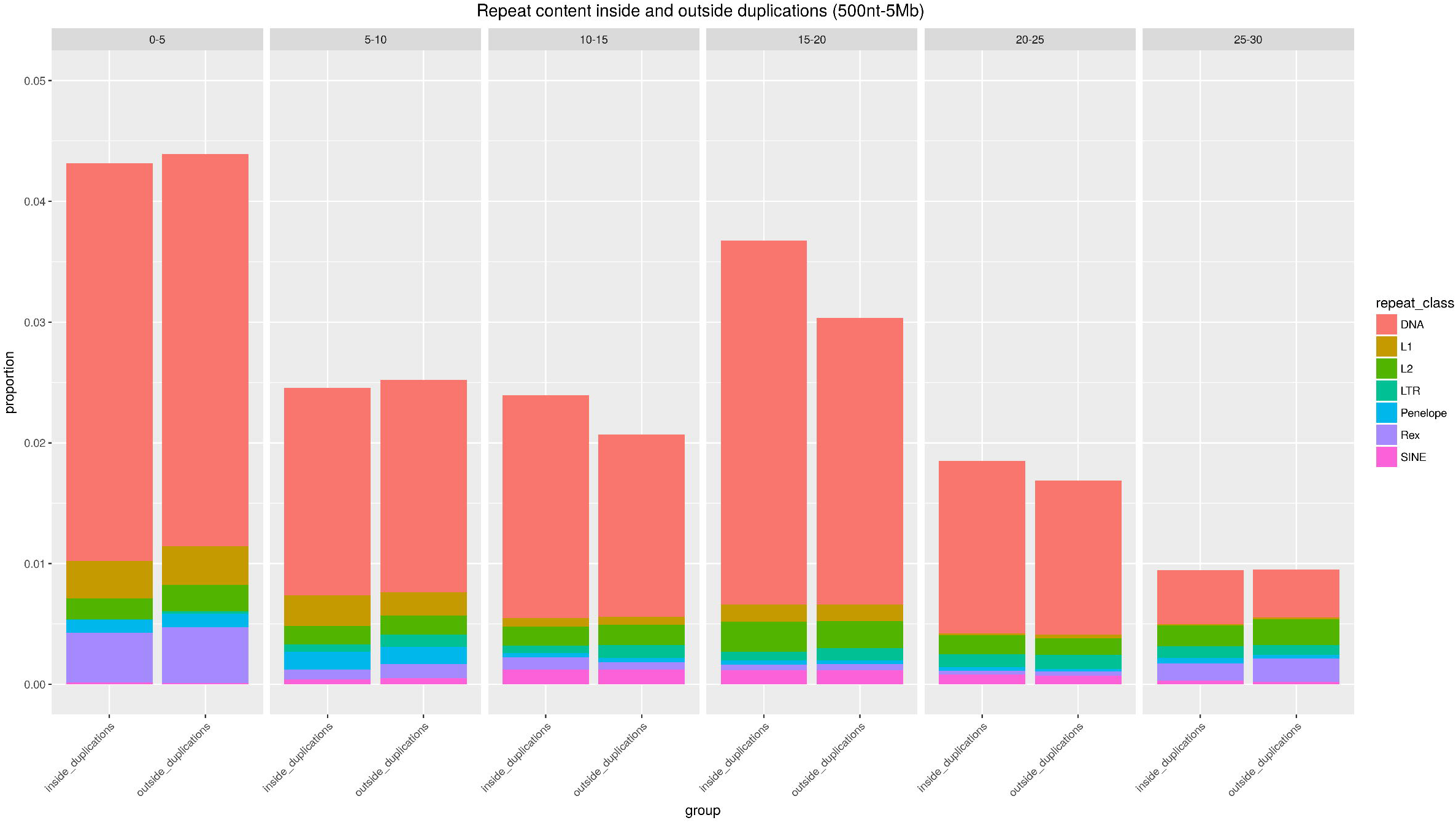
Proportion of nucleotides inside and outside *M. zebra* duplications which are part of a repeat element, grouped based on the percentage of divergence from the consensus. Colours and categories are defined as in Fig.3.

**Fig. 6.**
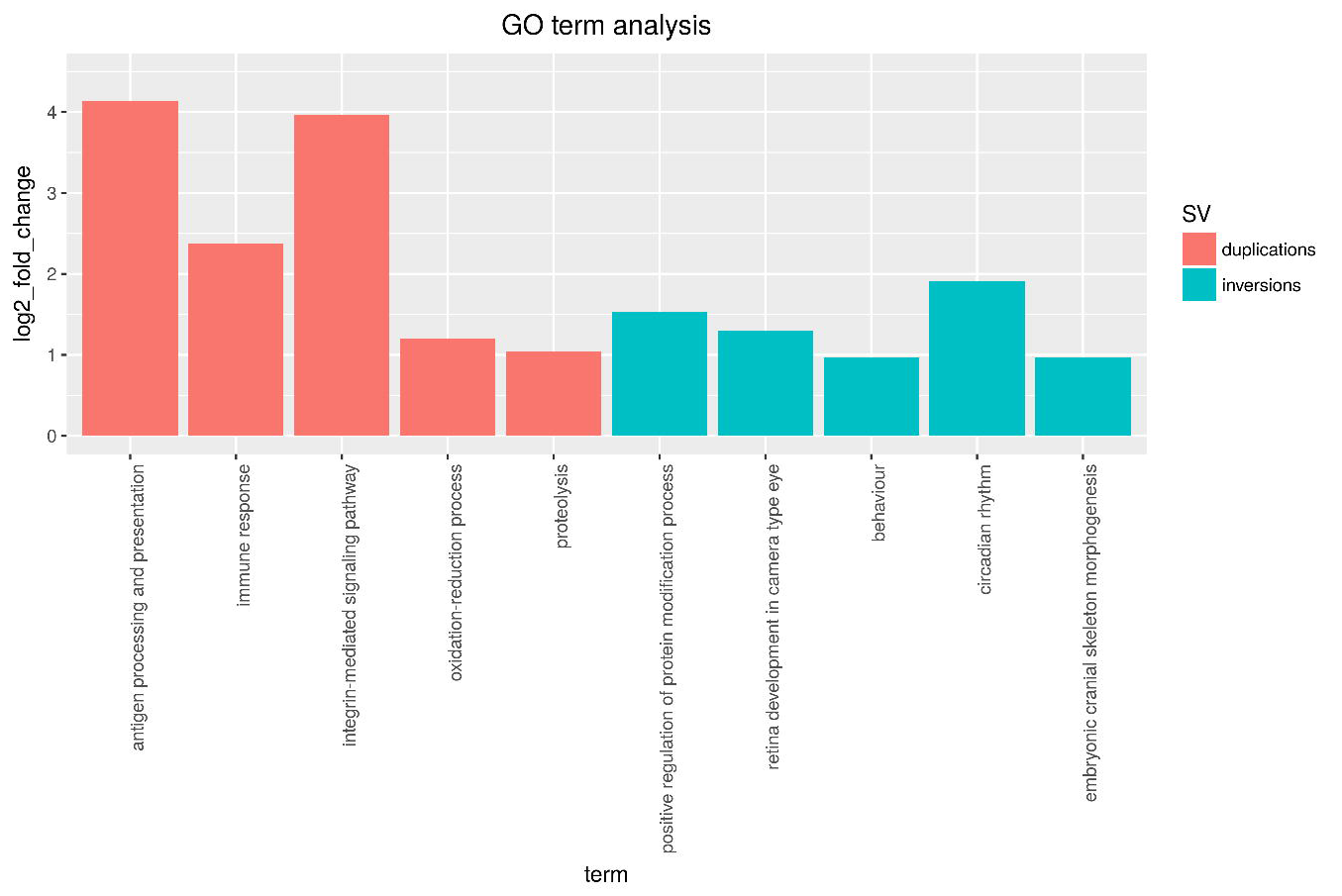
Selected GO terms found to be significantly enriched for the gene sets inside multi-species duplication and inversion (up to 5Mb) events.

**Fig. 7.**
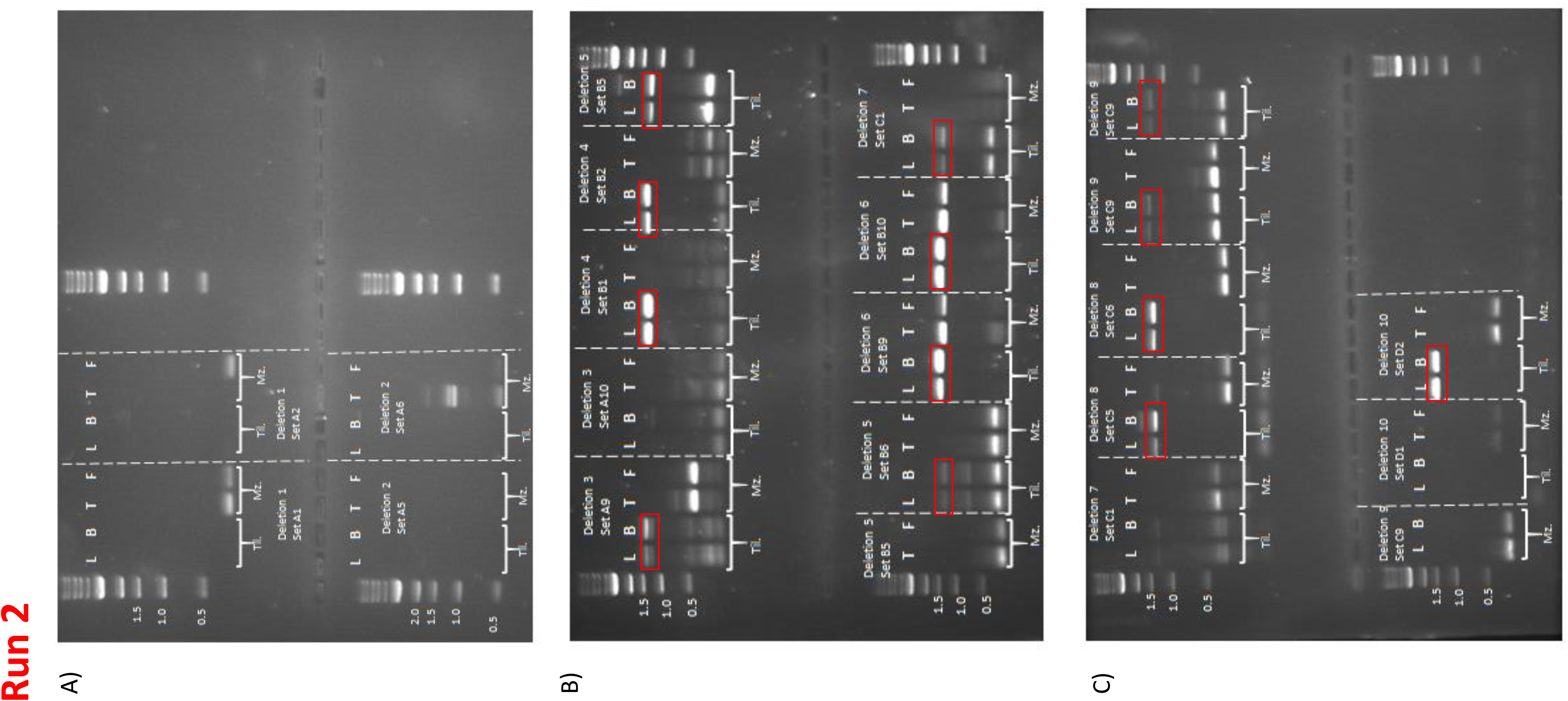
Gel images of PCR run 2 (validation of 10 deletion events in *M.zebra*). Red boxes indicate the expected product in the absence of the deletion (*O. niloticus* samples). No support was found for deletions 1,2 and 6. Key: L = liver, B = brain, T = Testis, F = fin, On = *O.niloticus*, Mz = *M. zebra*.

In order to identify homologous regions between the reference and each of the remaining species, we converted *Satsuma2* whole genome alignments to chain format, and performed a liftover of all SV coordinates from the reference to each of the other species. This allowed us to compare the repeat landscape in a pairwise fashion, considering different SV class separately.

When heterozygous, an inversion can favour the accumulation of mutations and novel transposable elements, as a result of reduced excisions rates (Charlesworth and Langley 1989; Charlesworth, et al. 1994). We tested this possibility by comparing the repeat content inside and outside inverted regions. We focused our analysis on the latest *O. niloticus* and *M. zebra* genome references, representing the highest quality assemblies among our five species. Overlap analyses based on the *O. niloticus* reference suggested a limited, but significant enrichment in DNA transposons inside inversions (size range: 500nt-5Mb; fold change=1.07, p=1e-04). While for other repeat classes the proportions are very similar inside and outside liftover inverted regions (Fig.3), the LTR representation is higher in the former across all divergence bins (Fig.4). This reflected in a significant enrichment in LTR elements inside inversions (size range: 500nt-5Mb; fold change=1.21, p=1e-04), as opposed to the LTR content outside inversions (fold change=0.92, p=1e-04).

Next, we repeated the analyses considering duplication events. We observed no difference in repeat content inside and outside duplicated regions (Fig.5). The association between duplications and LTR is weaker than expected (fold change=0.94, p=1e-04), while no significant deviation was found when considering DNA transposons (fold change=0.99, p=0.31). As for regions outside duplications, we observed a significant, although very limited, enrichment for both DNA (fold change=1.001, p=0.04) and LTR (fold change=1.05, p=8e-04) elements. It must be noted, however, that during the liftover conversion of the genomic coordinates, many inverted and duplicated regions were lost, limiting the sequence space considered in *M. zebra*.

### Enrichment of structural variants for developmental and immune related processes

Structural variation can provide important evolutionary novelty for speciation and the evolution of adaptive traits (Ohno 1970; Lynch 2002; Katju and Lynch 2006; Catacchio, et al. 2018). For instance, gene duplication can lead to dosage effects, neofunctionalisation or subfunctionalisation events (Lynch 2002; Katju and Lynch 2006), while inverted regions can experience drastically reduced recombination rates (Catacchio, et al. 2018). We took advantage of our SV dataset across 4 species, to investigate which genes are affected by duplication or inversion events.

We first considered different subsets of inversions, separating species specific events from the ones annotated in multiple species.

When looking at species specific events, we considered each species separately (Supplementary Table 1). We identified 559 genes in *H. burtoni*, 109 in *M. zebra*, 580 in *N. brichardi* and 814 in *P. nyererei*. Results for *H. burtoni* highlighted GO:0006955 (“immune response”, significant: 13; expected: 5.01; p_adj_ =0.0015), GO:0007600 (“sensory perception”, significant: 10; expected: 5.04, p_adj_ =0.03).

Inverted genes in *N. brichardi*, are enriched for GO:0065007 (“biological regulation”, significant: 193; expected: 154.35, p_adj_ =0.0411) and GO:0009416 (“response to light stimulus”, significant: 6; expected: 2.45, p_adj_ =0.0351). Interestingly, we also found one gene (*gja3*, coding for an intercellular channel) annotated to GO:0048050, (“post-embryonic eye morphogenesis”). *P. nyererei* genes are enriched for GO:0007602 (“phototransduction”, significant: 6, expected: 1.44, p_adj_ =0.003). Interestingly, this set also includes 2 genes annotated to GO:0002089 (“lens morphogenesis in camera-type eye”), *fn1a* and *foxe3*, as well as 3 genes annotated to GO:0061035 (“regulation of cartilage development”): *sox32*, *s1pr2* and *pthlha*. Members of the *sox* gene family encode for transcription factors, and play a crucial role in morphological and behavioural variation in teleosts (Voldoire, et al. 2017). *pthlha* is an oral jaw specific gene (Hulsey, et al. 2016) coding for the parathyroid hormone. In the case, of *M. zebra*, we could only identify one accession represented by 5 or more genes: GO:0006468 (“protein phosphorylation”; significant: 6; expected: 2.52;; p_adj_ =0.0374).

We also considered inversions which are common to at least 2 species, again not exceeding 5Mb in size (Supplementary Table 2). A total of 854 GO annotated genes (Supplementary Table 1) could be identified inside these SV regions. GO:term enrichment on this gene set highlighted accessions GO:0060041 (“retina development in camera-type eye”, significant: 14; expected: 5.7; p_adj_ =0.001), GO:0060042 (“retina morphogenesis in camera-type eye”, significant: 7; expected: 2.76, p_adj_ =0.0018), GO:0048706 (“embryonic skeletal system development”, significant 10; expected: 5.12; p_adj_ =0.03). Among the genes annotated to GO:0060041, we found: *vax2* (eye-specific homeobox gene), a gene known to regulate cone opsin expression (Alfano, et al. 2011); *fgf8a* (fibroblast growth factor 8a), part of a key pathway in animal evolution (Bloomquist, et al. 2017) and *ift172* (intraflagellar transport 172).

**Table 2.** Genomic coordinates and expected product size of the 10 tested deletions. Green and red colours indicate the events which could and could not be confirmed by PCR, respectively.

| <u>ID</u> | <u>genomic coordinates</u> | <u>Expected product size (kb): primer set 1 & 2</u> |
| --- | --- | --- |
| DEL_1 | contig401:86756-90467 | <u>Set 1</u>: 3.3, <u>Set 2</u>: 3.3 |
| DEL_2 | contig429:64638-70314 | <u>Set 1</u>: 5.2, <u>Set 2</u>: 5.2 |
| DEL_3 | lg1:9504350-9506050 | <u>Set 1</u>: 1.3, <u>Set 2</u>: 1.2 |
| DEL_4 | lg5:529562-531357 | <u>Set 1</u>: 1.3 <u>Set 2</u>: 1.4 |
| DEL_5 | lg11:29218518-29220314 | <u>Set 1</u>: 1.3 <u>Set 2</u>: 1.3 |
| DEL_6 | lg12:27848949-27850738 | <u>Set 1</u>: 1.4 <u>Set 2</u>: 1.4 |
| DEL_7 | lg16:40913013-40914775 | <u>Set 1</u>: 1.3 <u>Set 2</u>: 1.3 |
| DEL_8 | lg17:27332603-27334390 | <u>Set 1</u>: 1.3 <u>Set 2</u>: 1.3 |
| DEL_9 | lg18:23620664-23622449 | <u>Set 1</u>: 1.3 <u>Set 2</u>: 1.3 |
| DEL_10 | lg20:8700540-8702339 | <u>Set 1</u>: 1.3 <u>Set 2</u>: 1.3 |

We repeated the same procedure for inversions up to 10Mb in length, which increased the number of genes considered to 1,404 (Supplementary Table 2). While accession GO:0060041 was still significantly over-represented, we observed additional, immune related processes: GO:0019882 (“antigen processing and presentation”, significant: 8; expected: 2.7; p_adj_ =0.0044), GO:0006955 (“immune response”, significant: 19; expected: 12; p_adj_ =0.03) and GO:0042445 (“hormone metabolic process”, significant:6;expected: 2.14; p_adj_ =0.01). As part of GO:0006955, we found the gene *nfil3* (Nuclear Factor, Interleukin 3 Regulated), coding for a transcriptional regulator.

Next, we compared the GO terms across different subtrees of our five-species phylogeny (Supplementary Fig.2, Supplementary Table 3). We first identified inversions common to *M. zebra* and *P. nyererei* but absent in the other species, for which we could identify 210 inverted genes. We found enrichment for protein modification and processing, including GO:0006508 (“proteolysis”, significant: 14; expected: 7.3; p_adj_=0.014). When considering the branch leading to these two species as well as *H. burtoni* (134 genes, events absent in *N. brichardi*), we identified genes involved developmental processes, including 4 annotated to GO:0048598 (“embryonic morphogenesis”) and 2 genes for accession GO:0033339 (“pectoral fin development”): *cyp26c1* (cytochrome P450, family 26, subfamily C, polypeptide 1) and *sall4* (Spalt Like Transcription Factor 4). The former lies in a sex associated region in *H. burtoni* (Roberts, et al. 2016). Together with the fact that the inversion is lineage specific, it makes the gene particularly interesting. We can speculate that the inversion event might have helped the maintenance of specific haplotypes (including gene *cyp26c1*) through the suppression of recombination in the affected region, possibly contributing to the divergence of sex-associated traits.

The enrichment for developmental processes was also observed for genes in conserved inversions (up to 5Mb in size, n=90), among which GO:0060042 (“retina morphogenesis in camera-type eye”, <5 genes) and GO:0048706 (“neuron development”, significant 5; expected: 1.3; p_adj_ =0.04) are particularly interesting.

We also looked for genes contained inside tandem duplications, and filtered the resulting set based on evidence of tandem repeat of at least 3 consecutive exons in the target genome assembly (see materials and method). When considering species-specific events (Supplementary Table 4), we identified 204 genes in *H. burtoni*, 197 in *M. zebra*, 143 in *N. brichardi* and 224 in *P. nyererei*. For the *H. burtoni* gene set we identified, among others, GO:0006508 (“proteolysis”, significant: 19; expected: 7.3; p_adj_=0.021) and GO:0060078 (“regulation of postsynaptic membrane potential”,significant: 6; expected: 0.87; p_adj_=0.0002). Duplicated genes in *P. nyererei* are enriched for immune related processes, including GO:0006955 (“immune_response”, significant: 10, expected: 1.83; p_adj_=1.4e-05) and GO:0019882 (“antigen processing and presentation”, significant: 6; expected: 0.41, p_adj_<0.0001). Additionally, GO:0055085 is represented by 18 genes (“transmembrane_transport”, significant: 18; expected: 11.18, p_adj_ 0.028).

By requiring the duplication event to be shared by at least 2 species, we could identify 152 genes (Supplementary Table 4). Results highlighted the presence of GO:0019882 (“antigen processing and presentation”, p_adj_=1e^-6^), GO:0007229 (“integrin-mediated signalling pathway”, p_adj_=1.6e^-5^), and GO:0006955 (“immune response”, significant: 8; expected:1.54; p_adj_=1.5e^-4^). This dataset contains two genes encoding for an H-2 class II histocompatibility antigen chain: ENSONIG00000019943 and ENSONIG00000003904. Accession GO:0048854 (“brain morphogenesis”, significant:2; expected:0.15; p_adj_=1e^-3^) was also significant enriched, however it is represented by only 2 genes: *atp1a1* (ENSONIG00000012456), encoding for ATPase Na+/K+ transporting subunit alpha 1a, and *shank3* (SH3 and multiple ankyrin repeat domains 3). Similar to inversions, we looked at GO enrichment across the phylogenetic tree. While only one gene (*lyz*) was found in conserved duplications (after filtering for evidence of tandem repeats and presence in the *Ensembl* annotation), we had 41 genes for the *M. zebra*-*P. nyererei* subtree and 58 for the lineage including *H. burtoni* as well (Supplementary Fig.3). However, in all of these cases the significantly enriched terms were represented by very low (<4) numbers of genes.

Altogether, our analyses provide the first insights into the possible contribution of SVs to the evolution of adaptive traits in African Cichlids, including circadian rhythm, developmental processes and immune response mechanisms.

### Validation of selected deletion events

In order to better understand the reliability of our computational analyses, we decided to validate selected deletions by PCR amplification of the rearranged genomic region (Fig.7, Supplementary Fig. 4, Materials and Methods). We focused on 10, medium sized (1-5kb) deletion events annotated in *M. zebra* (Table 2). For the validation, we compared experimental results obtained using tissue samples for *M. zebra* (liver and brain) and *O. niloticus* (testis and fin). In this comparison, *O. niloticus* represented the SV-free reference sequence, while *M. zebra* is predicted to carry the deletion event (and hence show a smaller amplification product). Fig. 7 provides an overview of the results of the second PCR run. We could confidently confirm the deletion event in 7 out of 10 cases. In the case of deletions 1 and 2 of Fig.12, we were not able to detect the expected products. As for deletion 6, we had discordant results between run 1 (Supplementary Fig.4) supporting the SV, and run 2 showing the expected product in both *M. zebra* and *O. niloticus*. Even excluding deletion 6, we obtain a 70% concordance between our computational predictions and the PCR validation, providing evidence for the reliability of our bioinformatics pipeline.

## Conclusions

In this study, we provide an overview on the repertoires of structural variation in East African Cichlids, their evolutionary patterns, and their possible contribution to adaptation. We inferred the gain and loss patterns of all annotated SVs across the phylogenetic tree. This allowed us to separate lineage specific and more conserved events, and highlight high proportions of lineage specific gains, consistent across SV types. While the size distributions are generally comparable across different conservation levels, we see a shift towards larger sizes in the case of deeply conserved inversions.

We investigated the repertoires of genes affected by inversions and duplications, considering species specific and more conserved events separately. Our results highlight, for the first time, the possibility that large scale genomic rearrangements have played a crucial role in the adaptive radiation of East African Cichlids. Among the most interesting biological processes associated with inversions, we find “behavior” (GO:007610), “retina development in camera-type eye” (GO:0060041), “pectoral fin development” (GO:0033339) and “embryonic skeletal system development” (GO:0048706). Moreover, we found enrichment for “neuron development” (GO:0048666) associated with events in the Haplochromine lineage (*M. zebra*, *P. nyererei*, *H. burtoni*). GO:0033339 includes gene *Cyp26c1*, lying in a sex associated region identified in *H. burtoni* (Roberts, et al. 2016). This result opens to speculations on the link between inversion events, suppressed recombination, and the divergence of sex-associated traits across lineages.

When we consider duplicated regions, we find an enrichment for “antigen processing and presentation” (GO:0019882) and additional immune related categories. This “theme” is common to both species-specific and multi-species events. However, when we consider different subtrees separately, the number of significant genes drops dramatically, making us less confident about the biological relevance of the enrichment results. Our set of genes inside duplicated regions include an *H-2 class II histocompatibility antigen* locus, as well as *ilf2* (interleukin enhancer binding factor). The observed association between immune genes and duplication events is not surprising, being in line with previous studies on the fast, adaptive evolution of the vertebrate immune system (Bartl, et al. 1994; Das, et al. 2012; Lighten, et al. 2017). In fishes, differences in parasite communities across foraging habitats can determine strong selective pressure, favouring adaptive phenotypes and ecological speciation. In the threespine stickleback (*Gasterosteus aculeatus*), it has been shown that Major Histocompatability Complex (MHC) genes are linked with female mating preference, suggesting that divergent selection acting on MHC genes might influence speciation (Milinski, et al. 2005; Matthews, et al. 2010).

We investigated the possibility of differential evolutionary patterns between inverted and non-inverted regions by comparing their repetitive element landscapes in *M. zebra*. Despite the observation of a significant enrichment in both DNA transposons and LTR elements, we observed little difference in repeat content. This holds true when comparing duplicated regions to the rest of the genome, despite a slightly higher representation of highly diverged copies inside duplications.

Recently, Conte *et al.* generated an improved reference assembly for the Cichlids *M. zebra* and *niloticus* (Conte, et al. 2018). The authors compared the genome structure of the two species at the chromosome scale, taking advantage of the high quality of these references. They observed a high number of ~2-28Mb, intra-chromosomal SVs, but a limited number of inter-chromosomal rearrangements. They also identified structural changes associated with lower recombination rates, suggesting inversion events between the rock- and sand-dwelling species in Lake Malawi. Their analyses on repetitive elements indicate that *M. zebra* has a higher number of recent transposable element insertions compared to *O. niloticus*. While providing important insights into the rearrangement evolution of lake Malawi Cichlids, this study does not investigate the patterns of structural variation across the three main lakes of East Africa, or their possible contribution to speciation and adaptive phenotypes.

The evolution of Cichlids in African lakes represent a fascinating example of how relatively low degree of genetic variation can provide the substrate for an explosive and rapid radiation of species, allowing for the adaption to many different ecological niches. Single nucleotide variants, large scale rearrangements, transposable elements and several regulatory mechanisms can all play together, possibly resulting in powerful combinations of genetic traits with high adaptive potential. We are only starting to understand the evolutionary dynamics and molecular mechanisms underlying this impressive radiation, and much work is still needed to shed light on all different aspects and key players involved.

### Materials and Methods

Paired-end libraries available for *Neolamprologus brichardi*, *Metriaclima zebra, Pundamilia nyererei* and *Haplochromis burtoni* (Brawand, et al. 2014) were downloaded using *fastq-dump* from the *sra-toolkit* (https://www.ncbi.nlm.nih.gov/sra/docs/toolkitsoft/). Due to lower base quality issues, the last 30 nt at the 3’ end of the longest, 100nt reads were trimmed. All libraries were mapped against the *O. niloticus* genome assembly (Supplementary Fig.1) using *gmap* (Wu, et al. 2016). The resulting bam files were sorted and indexed using *samtools* (Li, et al. 2009), then used as input for 3 algorithms: *Breakdancer* (Fan, et al. 2014), *Delly* (Rausch, et al. 2012) and *Pindel* (Ye, et al. 2009).

SV predictions were first filtered by: a minimum of 2 libraries and 5 discordantly mapping read pairs supporting the call (*Breakdancer* and *Pindel*); a *Breakdancer* score of 99; both a PASSED and PRECISE flag provided by Delly’s output files. For each tool and rearrangement class separately (with the exception of translocations), we merged predictions with a reciprocal coordinate intersection of at least 90% into a single SV call.

The sets of merged filtered calls of each algorithms were then compared in a pairwise manner. Specifically, we used *Bedtools intersect* (Quinlan 2014) to identify SVs independently called by two different algorithms, with a reciprocal intersection at least 90% of the SV region. This gave us three sets of SVs supported by at least 2 algorithms: *Breakdancer*+*Delly, Breakdancer*+*Pindel* and *Delly*+*Pindel* (Supplementary Fig.5). The annotations of each SV class across all species was then combined into a single BED file. For each combined set, we then carried out a conditional merging of the SV genomic coordinates. For events up to 0.5kb in size, we required a minimum of 50% reciprocal intersection for multiple calls to be merged together. For size ranges of 0.5-1kb, 1-10kb and all events greater than 10kb, we used a threshold 80, 90 and 95%, respectively (in each range, we included the bottom value while excluding the top one).

### Overlap analyses and GO:term enrichment

Analyses of overlap between SVs and genome annotation were performed using *GAT* (Heger, et al. 2013). The *O_niloticus* UMD1 gene annotation was downloaded from the NCBI database (https://www.ncbi.nlm.nih.gov/genome/197?genome_assembly_id=391053) and converted into BED format. Genes inside SV regions were identified by comparing the gene annotation with the genomic coordinates of our SV dataset, using *Bedtools intersect* (Quinlan 2014). We selected genes fully contained inside an SV event by using the options “-a genes.bed -b SV.bed -f 1”.

We used a combination of *Biomart* (www.ensembl.org/biomart/martview/) and *DAVID* (https://david.ncifcrf.gov/) to map all NCBI gene ids to the corresponding Ensembl gene ids. GO:term enrichment was performed on the set of genes mapping to an Ensembl gene ids. We used the *elim* algorithm from the *R* package *TopGO* (Alexa, et al. 2006). The gene background was defined as the set of all genes in the NCBI annotation mapping to an *Ensembl* gene id.

**Table S1.** Results of MW test to compare SV size distribution across different conservation Categories. For each comparison, the p-value is indicated. In the case of a significant difference, the directionality of the change is indicated. For example, “1<2” indicates that the ranks of the 1-species dataset are significantly lower than those for the 2-species dataset.

| MW test p-value | 1 vs 2 | 2 vs 3 | 3 vs conserved |
| --- | --- | --- | --- |
| **DEL** | 0.38 | 0.29 | 0.0008 (3<cons) |
| **DUP** | 0.005 (1<2) | 0.008 (2<3) | 0.35 |
| **INV** | <0.0001 (1<2) | <0.0001 (2<3) | <0.0001 (3<cons) |

### Whole genome alignments and repeat content analyses

In order to generate whole genome alignments between the latest *M. zebra* (Conte and Kocher 2015) and *O. niloticus* (Conte, et al. 2017) assemblies, we ran *Satsuma2* (https://github.com/bioinfologics/satsuma2) using the following parameters:

~~~
-slaves 10
-threads 16
-km_mem 120
-sl_mem 120
-prob_table true
-min_prob 0.99999
-min_seed_length 20
-max_seed_kmer_freq 1
-min_matches 10
-dump_cycle_matches 1
~~~

In order to compare *Satsuma2* results with our dataset of *M. zebra* deletions (*O. niloticus* genomic coordinates), we converted the *satsuma_summary.chained.out* output file into a 6 columns BED file. We then used the command *bedtools intersect* from *Bedtools* (Quinlan 2014) to identify alignments of sequences overlapping a deletion event. A deletion was considered to be discordant with *Satsuma2* if at least one alignment spanning 50% or more of the predicted deleted region could be identified.

For the analysis of the repeat content inside and outside SV regions, we used the Cambridge Cichlid Browser (http://em-x1.gurdon.cam.ac.uk/cgi-bin/hgLiftOver) to liftover coordinates from *O. niloticus* to *M. zebra*, using the latest high quality PacBio assemblies. *RepeatMasker* was run on the *M. zebra* to identify repetitive elements genome-wide. The .out result file of *RepeatMasker* was reformatted to generate a 6 column BED file. For each SV separately, we then used *Bedtools intersect* (Quinlan 2014) to identify repeat elements fully contained inside the liftover SV coordinates. Overlapping repeat elements were separated based on the percentage of divergence from the consensus sequence, provided in the .out result file. An equivalent approach was used to identify repeat elements outside SV events. The repeat content was then calculated as the proportion of repeat nucleotide positions over the total length of the genomic space considered (total size of SV space or the genomic space outside SV).

### PCR validation of structural variants

Oligonucleotide primers were designed against the latest *O. niloticus* assembly using *Primer3* software (Untergasser, et al. 2012) and all primers were synthesised by Integrated DNA Technologies, Iowa.

DNA was extracted from samples of frozen tissue or tissues preserved in ethanol (25 mg) from *Maylandia zebra* (fin, testis) and *Oreochromis niloticus* (brain, liver), using the MagAttract HMW DNA Kit (Qiagen, CA) according to the manufacturer’s protocol for “Manual Purification of High Molecular Weight Genomic DNA from Fresh or Frozen Tissue”. Final DNA concentrations were determined using Qubit fluorometer™ 2.0 (Invitrogen, Life Technologies) and purity was assessed using A260:280 ratio (≥1.8) by measurement on a Nanodrop™ spectrophotometer (Thermofisher Scientific). PCR products were amplified according to manufacturer’s protocol for NEBNext® High-Fidelity 2X PCR Master Mix in 25µl reactions using 50 ng of DNA template. PCR cycle conditions were followed as stated in manufacturer’s protocol and extension times were adjusted according to length of expected product size. PCR products were visualised on 1.5% (w/v) agarose gels stained with SYBR™ Safe DNA Gel Stain and imaged using the Alliance 2.7 gel documentation system (UVITEC, Cambridge).

## Supporting information

## Supplementary Figures

Supplementary Fig. 1 Schematic of the SV detection pipeline

Supplementary Fig. 2 Association of different enriched GO terms across the phylogenetic tree, considering the genes found inside inverted regions (up to 5Mb). For each node, selected GO terms are shown for the inversion events specific to (and conserved across) the *M.zebra+P.nyererei* lineage (top right), *M.zebra+P.nyererei+ H.burtoni* lineage (top-centre) and conserved across all species (bottom left). Numbers on the top of each bar indicate the number of observed genes.

Supplementary Fig. 3 Association of different enriched GO terms across the phylogenetic tree, considering the genes found inside duplicated regions. For each node, selected GO terms are shown for the duplication events specific to (and conserved across) the *M.zebra+P.nyererei* lineage (top right), the *M.zebra+P.nyererei+H.burtoni* lineage (top-centre) and conserved across all four species (bottom left). Numbers on the top of each bar indicate the number of observed genes.

Supplementary Fig. 4 A) Experimental design for the PCR validation of deletion events. Arrows represent primer sequences mapped to the genomic sequence (in blue and red). Primer couple AF1 + AR1 is used to test for the presence or absence of the deletion event (expected to differ by about *N* bp in the amplification product). Primer couples BF1 + BR1 and CF1 + CR1 are used as a control (expected product:300-400bp). B-E gel images of PCR run 1, used for the validation of 10 deletion events. See Fig. 7 for a detailed explanation of the figure labels.

Supplementary Fig. 5 Venn *diagram* depicting the intersection between filtered SV calls of all three tools (*Breakdancer*, *Delly* and *Pindel*). In the case of insertions, no intersection was found between *Breakdancer* and the other two tools.

## Supplementary Tables

Supplementary Table 1. Names and corresponding GO annotation for different subsets of genes inside species specific inverted regions

Supplementary Table 2. Names and corresponding GO annotation for different subsets of genes inside inverted regions, considering events found in 2 or more species

Supplementary Table 3. Names and corresponding GO annotation for different subsets of genes inside inverted regions, considering events conserved across different branches of the phylogenetic tree.

Supplementary Table 4. Names and corresponding GO annotation for different subsets of genes inside species specific duplicated regions

Supplementary Table 5. Names and corresponding GO annotation for different subsets of genes inside duplicated regions (found in 2 or more species)

