## Supplementary figures and images for "Analysis of structural variants in four African Cichlids highlights an association with developmental and immune related genes"

### Supplementary file 1

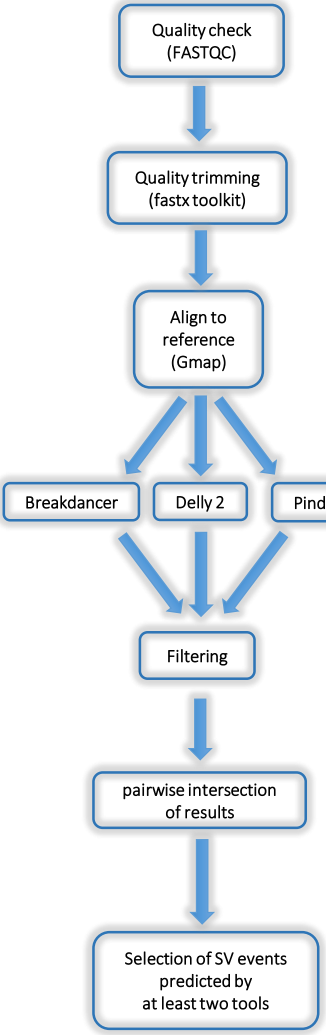

### Supplementary file 2

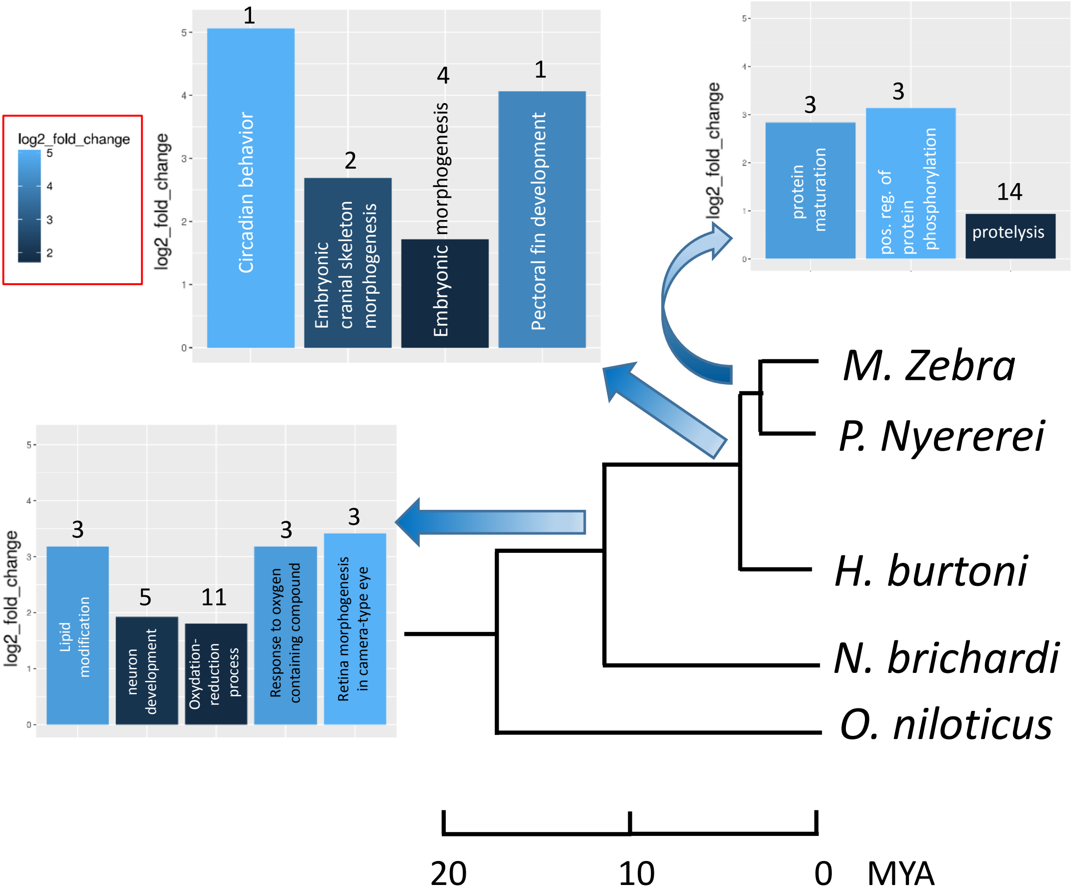

### Supplementary file 3

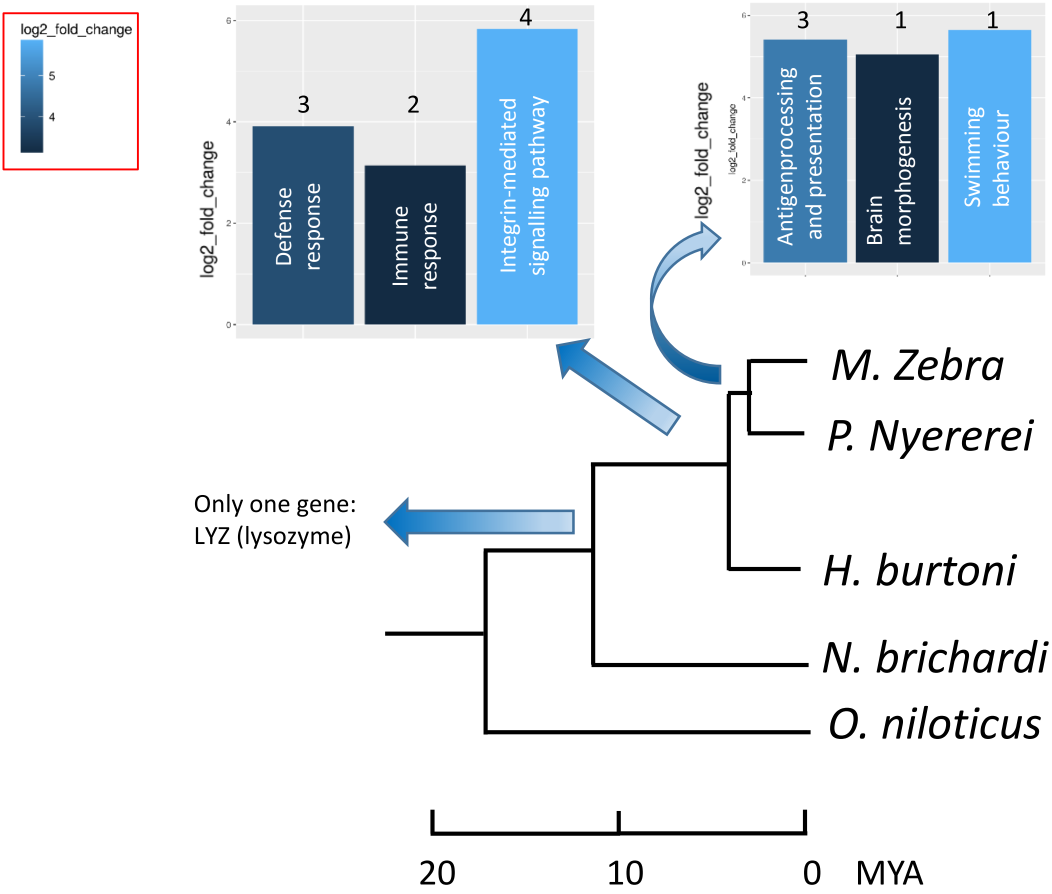

### Supplementary file 4

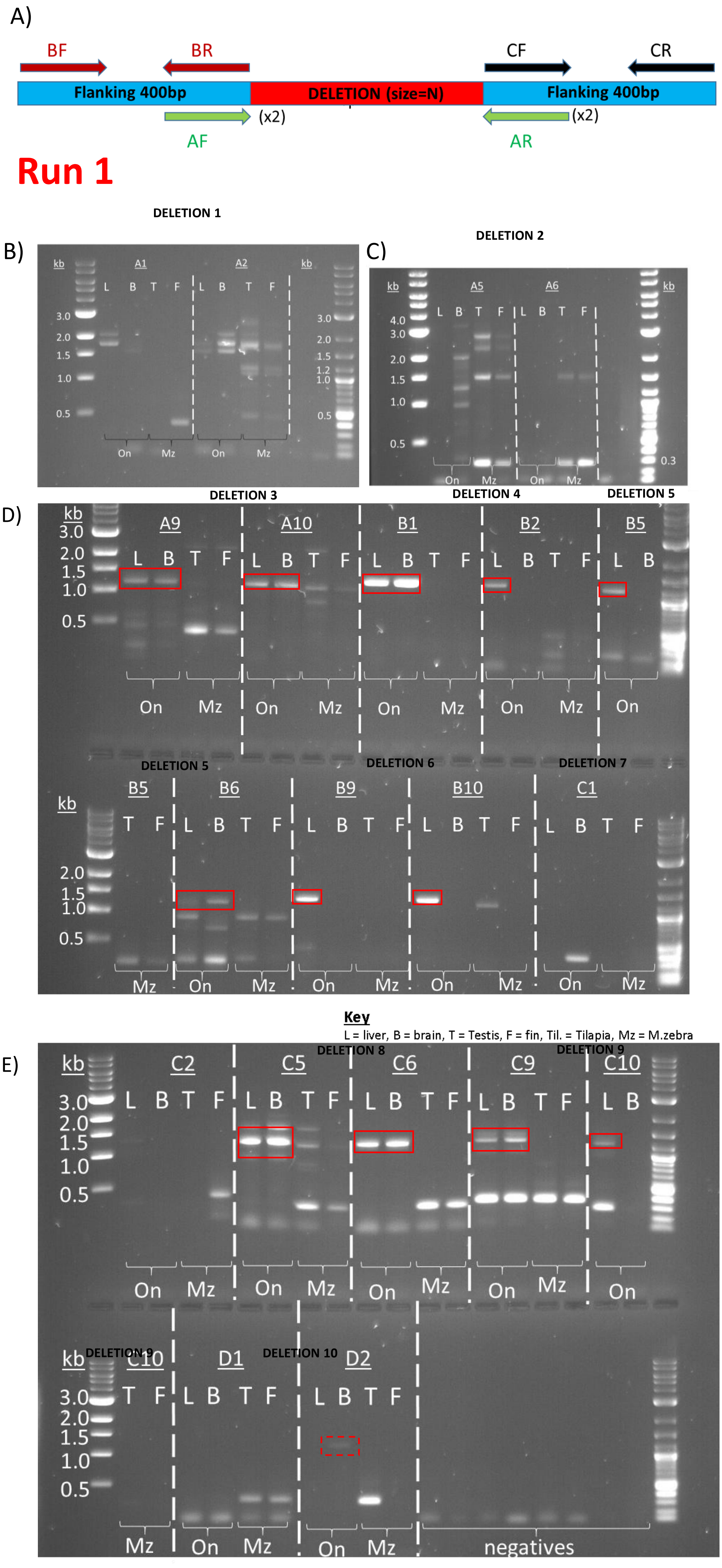

### Supplementary file 5

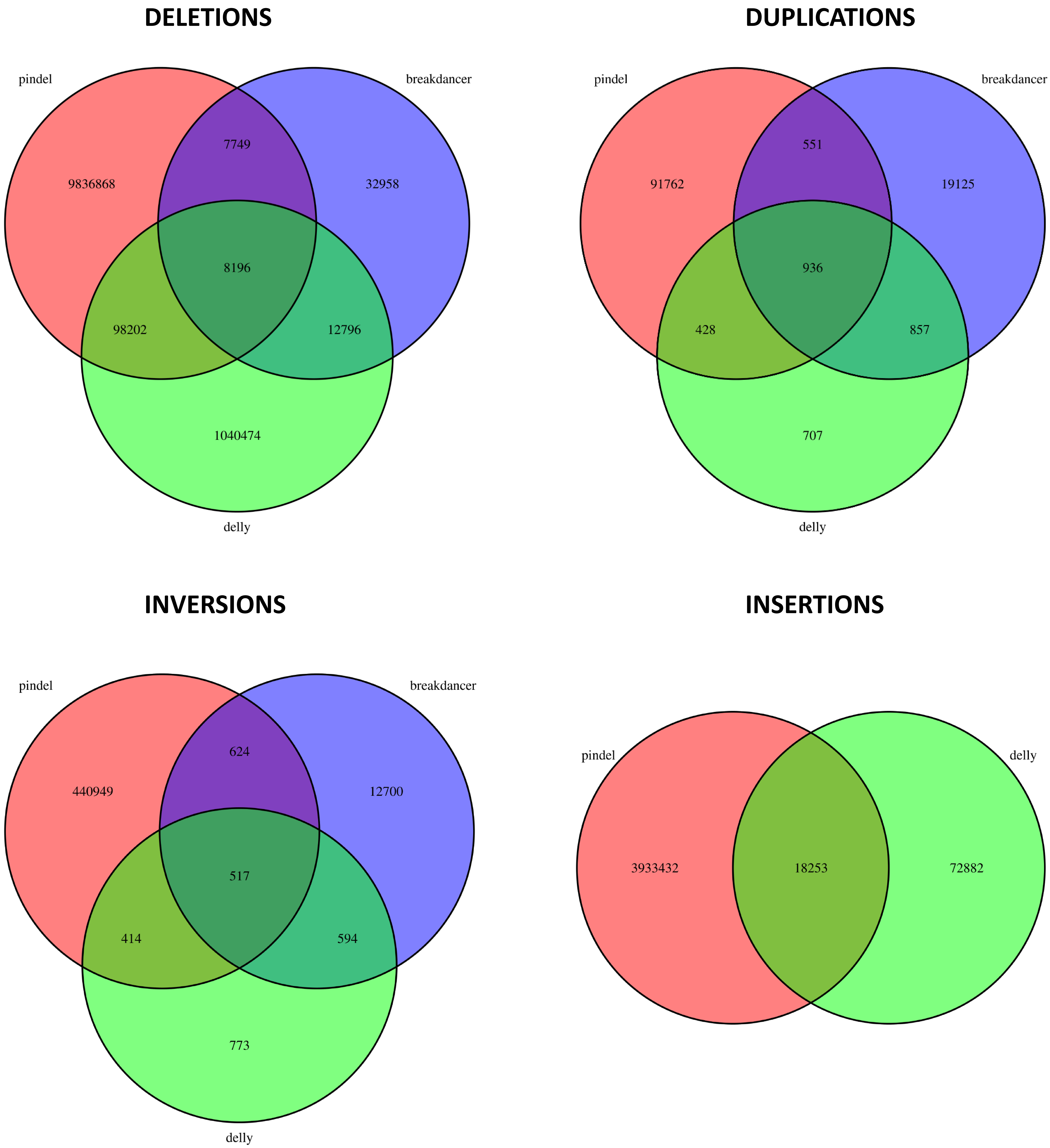
